# VIPER-seq: Protein-mediated small RNA enrichment enhances plant virus detection at low viral abundance

**DOI:** 10.64898/2026.10.26.761846

**Authors:** Marie-Emilie A. Gauthier, Flávia Sarti Bonora, Paul Richard Campbell, Roberto A. Barrero, Felipe F. de Felippes

## Abstract

Early and accurate detection of viral infections is critical for effective disease management and biosecurity programs. High-throughput sequencing technologies have emerged as powerful tools for plant disease diagnostics. In particular, sequencing of virus-derived small RNAs (vsiRNAs) enables the simultaneous detection of both known and novel viral agents. Compared with other approaches, small RNA sequencing (sRNA-seq) can offer superior sensitivity because it is less affected by the abundance of host messenger and ribosomal RNAs (mRNA and rRNA, respectively), and because sRNAs are generally more stable than longer RNA molecules. However, the sensitivity of sRNA-seq can be compromised by the presence of abundant endogenous small RNAs, which dilute the viral signal and hinder pathogen detection, particularly in low-titre infections. Here, we present VIPER-seq (Viral small RNA Isolation by Protein-mediated Enrichment and Recovery sequencing), a method that exploits sRNA-binding proteins to enrich vsiRNAs prior to high-throughput sequencing. Using cucumber seeds infected with cucumber green mottle mosaic virus (CGMMV), we demonstrate that VIPER-seq substantially enhances the detection of infected plant material compared with conventional sRNA-seq. The increased sensitivity is particularly evident under low-titre infection conditions, demonstrating the utility of VIPER-seq for the early detection of viral infections and its potential to strengthen plant disease diagnostics, seed health testing, and biosecurity surveillance programs.

## Introduction

Diseases caused by plant viruses are among the most significant challenges facing agriculture worldwide, causing annual losses estimated in the billions of dollars (Jones and Naidu, 2019). In severe cases, viral outbreaks can devastate entire plantations, reducing crop yields, affecting farmers’ livelihoods, increasing food prices, and, in extreme circumstances, threatening food security (Prasad *et al*., 2020; Sevik and Arli-Sokmen, 2012). Many of the crops most severely impacted by viral diseases are also major staple foods. Consequently, plant virus epidemics can have profound socioeconomic impacts and may contribute to food shortages among vulnerable populations that rely on these crops for subsistence (Jones and Naidu, 2019; Rybicki, 2015).

In an increasingly interconnected world, plant viruses can spread rapidly through both direct and indirect human activities, including the movement of plant material and agricultural trade (Ranawaka *et al*., 2020). Early detection of infected material is therefore critical for preventing disease spread, and accurate virus identification is a key prerequisite for implementing effective management and containment strategies (Rodoni, 2009). High-throughput sequencing (HTS) technologies have emerged as powerful and unbiased diagnostic tools capable of detecting a broad range of plant pathogens in a single assay (Massart *et al*., 2022). Among these, small RNA sequencing (sRNA-seq) has become an important approach for virus detection. During infection, plants mount an antiviral RNA-silencing response that results in the production of large quantities of virus-derived small interfering RNAs (vsiRNAs) (Csorba *et al*., 2015; Zhang *et al*., 2015). Because vsiRNAs are generated directly from viral genomes, they provide a rich source of diagnostic information for the detection and identification of viral pathogens.

Despite its advantages, the detection of low-titre infections by sRNA-seq can be challenging. Plant tissues contain large populations of endogenous short RNAs that are similar in size to vsiRNAs and can dominate sequencing libraries, thereby diluting the viral signal and limiting detection sensitivity. Consequently, an enrichment step that selectively increases the proportion of vsiRNAs prior to library preparation and sequencing could substantially improve the sensitivity of sRNA-seq-based diagnostics. Such an approach would not only enhance virus detection, particularly in low-titre infections, but also increase the cost-effectiveness of the method by reducing the sequencing depth required to obtain sufficient viral reads.

Viral suppressors of RNA silencing (VSRs) are a class of virus-encoded proteins that counteract the antiviral activity of vsiRNAs produced by the host plant (Csorba *et al*., 2015; Danielson and Pezacki, 2013). Several VSRs, including P15, P19, and P21, suppress RNA silencing by specifically binding small RNAs and preventing their incorporation into ARGONAUTE (AGO) proteins, the central effectors of the gene-silencing pathway (Chapman *et al*., 2004; Lakatos *et al*., 2004; Merai *et al*., 2006; Silhavy *et al*., 2002; Vargason *et al*., 2003; Ye *et al*., 2003). Owing to this highly specific RNA-binding activity, VSRs have become valuable tools for elucidating the molecular mechanisms underlying RNA silencing and, in biotechnology, for the detection of small RNAs and the enhancement and stabilisation of transgene expression in transient expression systems (Csorba *et al*., 2015; Danielson and Pezacki, 2013). Consequently, VSR-mediated capture of vsiRNAs represents an attractive strategy for enriching these molecules and enhancing the detection sensitivity of diagnostic approaches such as sRNA-seq.

Here, we present VIPER-seq (**V**iral small RNA **I**solation by **P**rotein-mediated **E**nrichment and **R**ecovery **seq**uencing), an sRNA-seq-based approach that uses the viral suppressor of RNA silencing P19 to capture and enrich vsiRNAs from infected plant tissues. Unlike conventional sRNA-seq workflows, which rely on total RNA as the starting material, VIPER-seq employs a streamlined protocol in which P19 immobilised on magnetic beads selectively captures vsiRNAs directly from plant tissue extracts. By increasing the proportion of viral small RNAs in sequencing libraries, VIPER-seq substantially enhances virus detection compared with conventional sRNA-seq, particularly when infected material is scarce or viral titres are low. These results demonstrate the potential of VIPER-seq as a highly sensitive and cost-effective tool for the detection of low-titre viral infections.

## Results

### VSRs can be used to isolate and enrich 21nt vsiRNAs for sRNA-seq analysis

We propose that VSRs could be used to capture vsiRNAs from infected plant material, thereby enriching these molecules for downstream sRNA-seq analysis (Figure 1A). To test this concept, hemagglutinin (HA)-tagged versions of the VSRs P19 and P15 (P19-HA and P15-HA, respectively) were co-expressed with an infectious clone of tobacco mosaic virus expressing GFP (TMV-GFP) (Lindbo, 2007) in *Nicotiana benthamiana* leaves by agroinfiltration. This experimental system enables rapid and controlled viral infection while maximising interactions between the VSRs and the vsiRNAs generated during the antiviral RNA-silencing response. Three days after infiltration, strong GFP fluorescence was detected in infiltrated leaves, indicating robust viral replication (Figure 1B). Following sample collection, a portion of the leaf tissue was used for total RNA extraction (P15_tot and P19_tot), which served as control samples. The remaining tissue was subjected to immunoprecipitation (IP) of the respective HA-tagged VSRs using anti-HA magnetic beads, followed by extraction of the associated RNAs (P15_IP and P19_IP). Both total RNA and VSR-bound RNA fractions were subsequently analysed by sRNA-seq (Figure 1C).

**Figure 1.**
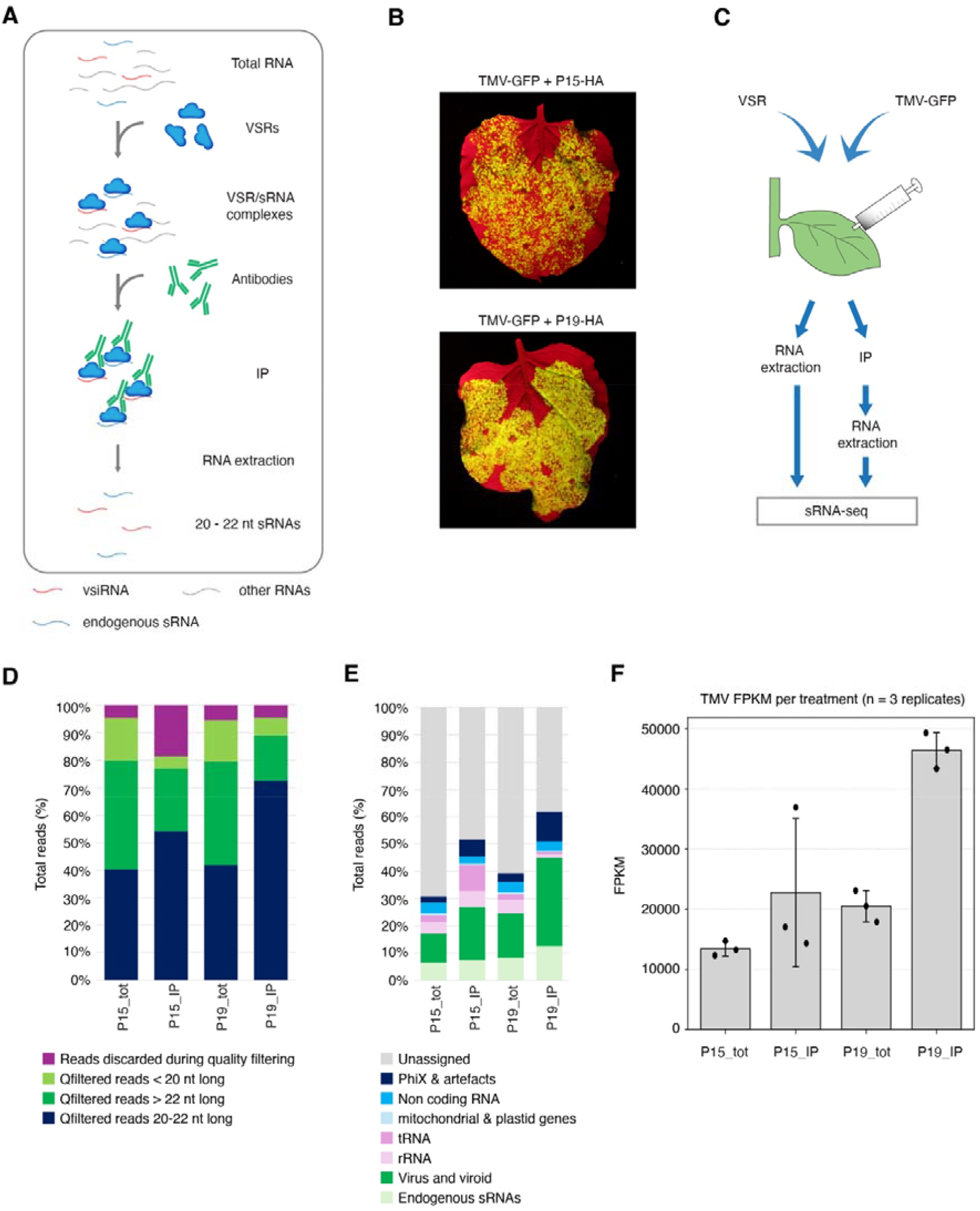
Proof-of-concept evaluation of VSR-mediated enrichment of vsiRNAs to improve sRNA-seq sensitivity. (A) Schematic overview of the approach. (B) *N. benthamiana* leaves three days after agroinfiltration with TMV-GFP and VSR constructs. (C) Experimental workflow used to assess vsiRNA enrichment. The proportion of sequencing reads by size class (D) and genomic origin (E) is shown. (F) Reads mapped to the TMV genome are presented as fragments per kilobase of transcript per million mapped reads (FPKM) across three biological replicates.

Previous characterisation of P15 and P19 binding affinities demonstrated that both proteins exhibit a strong preference for 20-22 nt sRNAs, functioning as molecular callipers that select sRNAs based on size (Merai *et al*., 2006; Vargason *et al*., 2003). Analysis of the size distribution of reads recovered from the IP fractions corroborated these findings. Reads in the P15_IP libraries were predominantly 21-22 nt in length, whereas P19_IP libraries were enriched for 20-22 nt sRNAs, with a pronounced peak at 21 nt (Figure 1D and Figure S1). In contrast, total RNA libraries contained a broader size distribution, consisting primarily of 20-24 nt sequences and a substantial proportion of reads shorter than 20 nt. As expected, the size-selective activity of these VSRs increased the relative abundance of vsiRNAs in the IP fractions and, consequently, the proportion of reads mapping to the viral genome (Figure 1E-F, Figure S1 and Table S1). Although P15-HA clearly improved sRNA-seq quality, its performance was less consistent than P19-HA, particularly in reproducibility and duplicate read levels. Consequently, all subsequent experiments were conducted exclusively with P19-HA.

In contrast to our standard experimental conditions, where the virus accumulated to high levels, virus detection often involves low-titre infections or only a few infected samples in a predominantly non-infected batch. We therefore investigated whether the beneficial effects of P19 on sRNA-seq performance are maintained under conditions of reduced viral abundance. To this end, we repeated the agroinfiltration assay using 1×, 10×, and 100× dilutions (1x_dil, 10x_dil, and 100x_dil, respectively) of the *Agrobacterium* culture carrying the infectious clone while keeping P19-HA levels constant (Figure S2A-B). Across all dilutions tested, P19-HA treatment resulted in a clear enrichment of 20-22 nt reads and a corresponding increase in vsiRNA abundance (Figure S2C-D and Figure S3D-E). Consistent with these observations, P19-HA-treated samples also exhibited improved read counts, FPKM values, and mean read depth relative to total RNA controls (Figure S2 and Figure S3). Notably, the percentage of the viral genome covered by at least 20 reads (PCT 20x) began to decline in total RNA samples expressing the lowest level of virus (100x_dil). This deterioration was less pronounced in the corresponding P19_IP samples (Figure S3C), suggesting that P19-mediated enrichment may be particularly advantageous for detecting viruses present at low titres.

### Purified, bead-immobilised P19 is an effective strategy to capture vsiRNAs from virus-infected plant extracts

The experimental design in our proof-of-concept study relied on co-expressing P19-HA and the TMV-GFP infectious clone within the same tissue. Although this approach effectively ensured interaction between P19-HA and vsiRNAs, it does not reflect real-world diagnostic scenarios, in which P19-HA is absent from the material being tested for viral infection. Therefore, for VIPER-seq to be broadly applicable, an efficient strategy is required to deliver P19-HA to the sample of interest and enable the capture of vsiRNAs.

Using *N. benthamiana* as a model system, we evaluated four different strategies for P19-HA delivery (Figure 2A). In Approach 1, leaf tissues independently expressing either P19-HA or TMV-GFP were combined to generate a mixed plant extract, which was subsequently subjected to P19-HA immunoprecipitation. In Approach 2, P19-HA expressed in *N. benthamiana* leaves was first purified and immobilised on magnetic beads. The P19-HA-coated beads were then incubated with plant extracts prepared from TMV-GFP-infected leaves. Approaches 3 and 4 followed the same principles as Approaches 1 and 2, respectively, except that total RNA isolated from TMV-GFP-infected tissue was used in place of infected plant extracts. Thus, in Approach 3, P19-HA-expressing tissue was mixed with total RNA from infected plants, whereas in Approach 4, purified bead-immobilised P19-HA was incubated directly with the RNA preparation.

**Figure 2.**
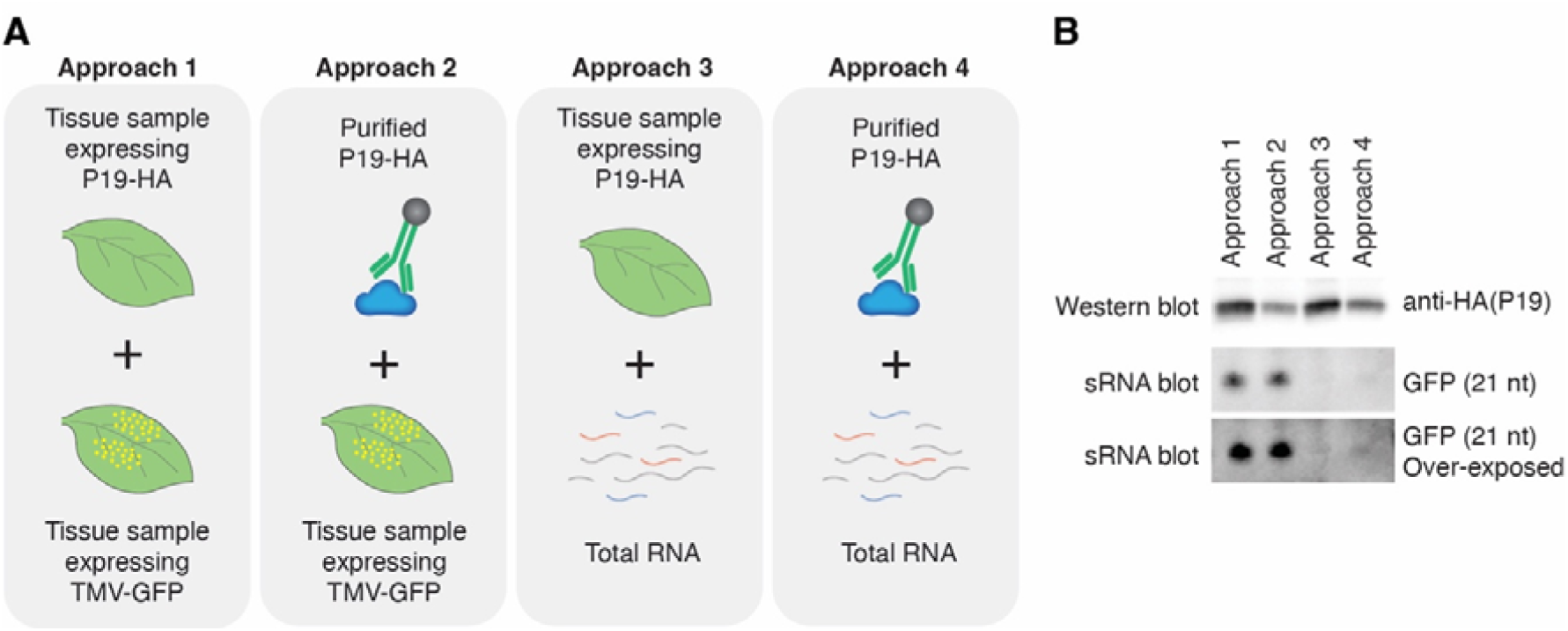
Assessment of different strategies for P19 delivery to infected plant tissue. (A) Four P19 delivery approaches were evaluated using the *N. benthamiana* infection system. (B) Western blot analysis with an HA-specific antibody confirmed the recovery of P19 following immunoprecipitation, while an sRNA Northern blot targeting GFP was used to assess the abundance of sRNAs co-immunoprecipitated with the VSR.

Northern blot analysis showed that vsiRNAs could be recovered using all four delivery strategies. However, substantially higher recovery efficiency was achieved when infected plant extracts, rather than purified RNA, were used as the source material (Approaches 1 and 2; Figure 2B). The lower efficiency observed with total RNA likely reflects the need for a more RNA-compatible workflow, including the use of RNase inhibitors and optimised conditions to preserve RNA integrity and facilitate P19-vsiRNA interactions. Nevertheless, we consider Approach 2 to be the most promising strategy because it offers greater flexibility regarding the source of the VSR and can be readily adapted to different sample types and experimental conditions. Consequently, this method was selected for P19-HA delivery in all subsequent experiments.

### VIPER-seq significantly increases virus detection in infected seeds

Having established the basis for VIPER-seq, we next assessed its performance relative to conventional sRNA-seq for virus detection in infected crop plants. Seeds represent an ideal material for this evaluation. They are the primary form of germplasm used for crop storage, exchange, and transport, and therefore are routinely screened for viral contamination before distribution to growers. In addition, RNA extraction from seeds presents several challenges that can compromise RNA quality and yield (Vennapusa *et al*., 2020), thereby reducing the effectiveness of RNA-seq-based analyses.

To evaluate VIPER-seq under these conditions, cucumber (*Cucumis sativus*) seeds infected with cucumber green mottle mosaic virus (CGMMV) were generated. Infection was confirmed by RT-qPCR detection of viral nucleic acids from individual seeds. Because virus testing is typically performed on seed batches rather than individual seeds, samples containing different proportions of infected and non-infected material (1:100, 1:400, and 1:800) were prepared by mixing ground infected and healthy seed tissue at the appropriate ratios. Each sample was then divided into two fractions. One fraction was used for total RNA extraction and subsequent conventional sRNA-seq analysis, while the second fraction was processed into a seed extract to which bead-immobilised P19-HA, previously purified from agroinfiltrated *N. benthamiana* leaves, was added for VIPER-seq analysis (Figure 3A).

**Figure 3.**
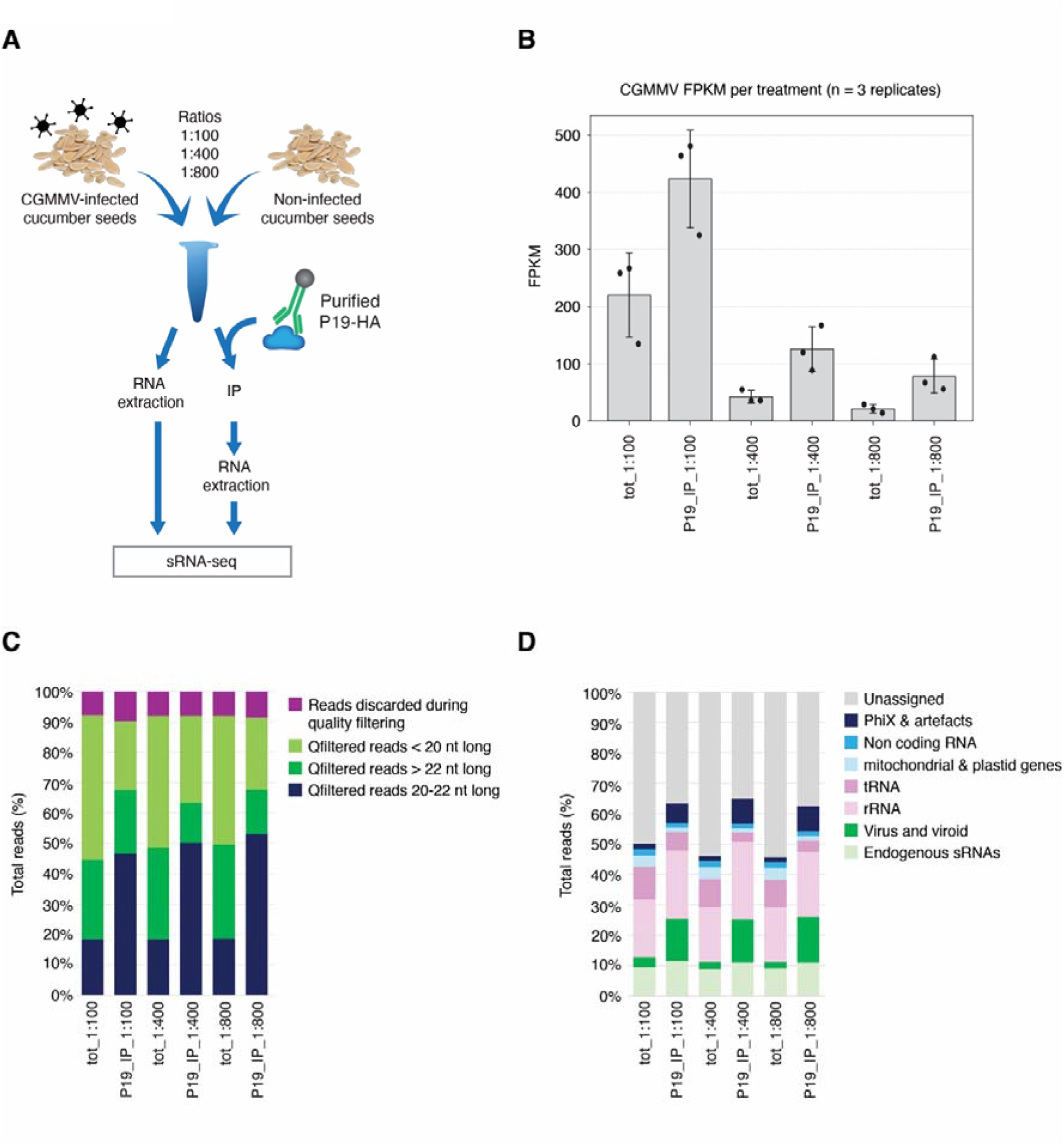
Efficiency of VIPER-seq for the detection of infected seeds. (A) Cucumber seeds infected with CGMMV were mixed with healthy seeds at three different ratios. Virus detection was performed using VIPER-seq by adding bead-immobilised P19-HA to crude plant extracts and was compared with conventional sRNA-seq performed on total RNA isolated from the same samples. (B) Sequencing reads mapping to the CGMMV genome are presented as fragments per kilobase of transcript per million mapped reads (FPKM) for three biological replicates. The distribution of sequencing reads across different size classes (C) and genomic origins (D) is shown.

Consistent with the results obtained using *N. benthamiana*, VIPER-seq outperformed conventional sRNA-seq for virus detection in seed samples. When the number of reads mapping to the viral genome was used as a measure of sensitivity, VIPER-seq achieved a 2- to 6.5-fold increase over standard sRNA-seq, with the greatest improvement observed in samples containing the lowest proportion of infected material (Figure 3B, Figure S4 and Table S1). This enhanced sensitivity, particularly in low-ratio samples, was accompanied by improved PCT20x values (Figure S4), further supporting our earlier observation that VIPER-seq is especially advantageous for the detection of low-titre infections and the analysis of large batches of test material. In accordance, VIPER-seq libraries were significantly enriched for 20-22 nt reads, leading to an increase in the proportion of vsiRNAs and depletion of non-informative sequences (Figure 3C-D and Figure S4).

## Discussion

The superior performance of VIPER-seq is largely attributable to the ability of P19 to selectively enrich vsiRNAs while excluding non-informative RNA species. This advantage becomes particularly evident in seed-derived samples, which contain abundant short RNA fragments that are likely generated through RNA degradation and high endogenous RNase activity (Figure 3). By preferentially capturing 20-22 nt vsiRNAs, P19 reduces background noise and increases the proportion of informative virus-derived sequences, thereby enhancing both the sensitivity and robustness of downstream virus detection. The benefits of P19-mediated enrichment may be further enhanced through protein engineering. P19 is highly amenable to structural modification, and several engineered variants have been shown to exhibit increased affinity for small RNAs. For example, linking two P19 monomers with a semi-rigid peptide linker increased sRNA-binding affinity by approximately 3.5-fold relative to the wild-type protein (Cheng *et al*., 2007). Similarly, substitution of a single amino acid residue (T111S) was sufficient to increase the affinity of P19 for the human miRNA miR122 (Cheng *et al*., 2011). These findings suggest that further optimisation of P19, whether through rational protein engineering or directed evolution approaches, could yield variants with enhanced affinity for vsiRNAs and, consequently, even greater sensitivity for VIPER-seq-based virus detection.

In this study, P19-HA was produced by agroinfiltration of *N. benthamiana* leaves, a highly attractive platform for the rapid and high-yield production of recombinant biological materials (Bally *et al*., 2018). Within only three days, large amounts of P19-HA-containing tissue can be generated and stored at -80°C, providing a stable and readily available long-term source of the VSR for VIPER-seq applications. It is important to note that, although the workflow incorporates high-salt washing steps designed to remove host-derived molecules associated with P19, residual *N. benthamiana* sequences may still be carried over into the final sequencing libraries. While such sequences can be readily identified and removed through standard bioinformatic filtering and are therefore unlikely to compromise virus detection, their presence may be undesirable in routine diagnostic laboratories and certified testing environments. Importantly, the use of *N. benthamiana* is not a prerequisite for VIPER-seq. P19 can be efficiently expressed in a range of heterologous systems without loss of functionality, including bacterial, human, and Drosophila expression platforms (Foss *et al*., 2019; Liu *et al*., 2012; Van Den Beek *et al*., 2014). The availability of these alternative production systems offers opportunities to generate highly purified P19 preparations that may be better suited to regulatory and diagnostic applications.

Early detection and accurate identification of plant viruses are critical for effective disease management and containment. sRNA-seq has become a valuable diagnostic tool, enabling the simultaneous detection and identification of multiple pathogens. In this study, we demonstrate that sequencing vsiRNAs captured by VSRs, rather than preparing libraries from total RNA, can substantially enhance the sensitivity of sRNA-seq. This improved method, here referred to as VIPER-seq, is thus a valuable new tool allowing for efficient virus detection from plant tissue.

## Material and methods

### Plant material

*N. benthamiana* plants were grown at 25 ºC, under long-day conditions (16:8 h light/dark photocycle). *C. sativus* ‘Green Gem’ seeds infected with CGMMV were generated by manual inoculation of CGMMV isolate Vir-5386 from the QDPI virus collection. Inoculation was performed 12 weeks post emergence, and tagged female flowers were manually pollenated with male flowers from other infected plants. Fruit were allowed to develop till yellow, seeds harvested fermented for two days and dried for storage. Cucumber “Long Green Supermarket” (Bunnings, Australia) was used as non-infected seed material.

### Plant transformation and GFP images

*Agrobacterium tumefaciens* strain GV3101was used for transient transformation of 4-week-old *N. benthamiana* leaves by agroinfiltration, as previously described (de Felippes and Weigel, 2010). For VSR expression, *A. tumefaciens* cultures were infiltrated at an OD_600_ of 0.4. For delivery of the TMV-GFP infectious clone, plants were infiltrated with *A. tumefaciens* cultures at OD_600_ values of 0.01 (1x_dil), 0.001 (10x_dil), or 0.0001 (100x_dil). Infiltrated leaves were harvested 3 days post-infiltration, and GFP fluorescence images were acquired as previously described (Gauthier *et al*., 2026).

### Molecular constructs

The coding sequences of P15 from peanut clump virus (PCV) and p19 from tomato bushy stunt virus (TBSV) were obtained from NCBI (RefSeq accession numbers NP_620041 and NP_062901.1, respectively). The sequences were synthesised as gBlocks (Integrate DNA Technologies, IDT, Coralville, USA) and designed to include a C-terminal HA tag, with XmaI and NotI restriction sites flanking the 5’ and 3’ ends, respectively. The VSR coding sequences were cloned into the pRBCS::GiFiP::tHSP vector (de Felippes *et al*., 2020) via the XmaI and NotI restriction sites, replacing the GiFiP coding sequence.

### RNA extraction

Total RNA was extracted from approximately 0.2 g of tissue ground in liquid nitrogen by addition of 1 mL TRIzol reagent (Thermo Fisher Scientific, Waltham, MA, USA). Following the addition of 0.2 mL chloroform, samples were centrifuged at 12,000 x g for 15 min at 4°C. The aqueous phase was transferred to a fresh tube and mixed with an equal volume of 100% ethanol. RNA was subsequently purified using the Direct-zol™ RNA MiniPrep Plus Kit (Zymo Research, Irvine, CA, USA) according to the manufacturer’s instructions. To generate samples containing different proportions of infected and non-infected RNA, independently extracted total RNA preparations were combined at the desired ratios.

RNA from immunoprecipitation (IP) fractions was extracted by adding TRIzol reagent directly to the resuspended beads following the final IP wash step and processing the samples according to the manufacturer’s protocol. To facilitate RNA precipitation and improve pellet visualisation, 2 μL of glycogen (20 mg/mL; molecular biology grade; Thermo Fisher Scientific, Waltham, MA, USA) was added during the isopropanol precipitation step.

### Immunoprecipitation assay

For experiments involving co-expression of the VSR and TMV-GFP, infiltrated *N. benthamiana* leaves were ground to a fine powder in liquid nitrogen. A 0.3 g tissue sample was resuspended in 1.5 mL of ice-cold IP buffer (50 mM Tris-HCl, pH 7.5; 150 mM NaCl; 10% glycerol; 0.1% Nonidet P-40) supplemented with cOmplete™ Mini EDTA-free Protease Inhibitor Cocktail (1 tablet per 10 mL; Roche, Mannheim, Germany), added immediately before use. Samples were incubated for 15 min at 4 °C with rotation and subsequently centrifuged at 4,000 × g for 5 min at 4 °C to remove cellular debris. The supernatant was filtered through Miracloth into a fresh tube. Immunoprecipitation of the VSR was performed by adding 15 μL of Pierce™ Anti-HA Magnetic Beads (Thermo Fisher Scientific, Waltham, MA, USA) to the extract, followed by incubation for 2 h at 4 °C with rotation. Beads were collected using a magnetic rack, washed three times with IP buffer, and finally resuspended in 100 μL of the same buffer.

The immunoprecipitation experiment involving cucumber seeds consisted of two stages. First, bead-immobilised P19-HA was purified from infiltrated *N. benthamiana* leaves (0.3 g tissue per sample) as described above, with the following modification. After the standard wash steps, beads were incubated for 5 min in high-salt wash buffer (50 mM Tris-HCl, pH 7.5; 1 M NaCl; 10% glycerol; 0.1% Nonidet P-40), followed by two additional washes with IP buffer. The beads were then resuspended in IP buffer supplemented with cOmplete™ Mini EDTA-free Protease Inhibitor Cocktail and stored at 4 °C until use. A volume of 30 μL of bead suspension was obtained from each 0.3 g of starting leaf material. In the second stage, CGMMV-infected and non-infected cucumber seeds were ground separately and subsequently combined at the desired ratios. Cell extracts were prepared by resuspending 0.3 g of tissue in 1.5 mL of IP buffer supplemented with cOmplete™ Mini EDTA-free Protease Inhibitor Cocktail, followed by incubation for 15 min at 4 °C with rotation. After centrifugation at 4,000 × g for 5 min at 4 °C and filtration through Miracloth, 30 μL of purified bead-immobilised P19-HA was added to each extract and incubated for 2 h at 4 °C with rotation. Beads were then washed three times with IP buffer and resuspended in 100 μL of the same buffer.

### P19 delivery test

All *N. benthamiana* cell extracts and IP procedures used in the P19 delivery assay were performed as described in the Immunoprecipitation Assay section. For Approach 1, IP was carried out using a cell extract prepared from equal amounts of liquid nitrogen-ground TMV-GFP- and P19-HA-expressing *N. benthamiana* leaf tissue (0.3 g of each tissue in 3 mL of IP buffer). Approach 2 involved the immunoprecipitation of P19-HA from *N. benthamiana* leaves, followed by the addition of bead-immobilised P19-HA to a cell extract prepared from TMV-GFP-infected tissue, as described for the cucumber seed immunoprecipitation assay. For Approaches 3 and 4, 6 μg of total RNA extracted from TMV-GFP-infected *N. benthamiana* leaves was incubated with either a cell extract prepared from P19-HA-expressing tissue or 30 μL of purified bead-immobilised P19-HA, respectively. All samples were incubated for 2 h at 4 °C with rotation, washed, resuspended in IP buffer, and processed for RNA extraction as described above.

### Northern and Western blot analysis

Northern blots for sRNA detection were done by running all RNA recovered from the IP procedure on a 17% polyacrylamide electrophoresis gel containing 7 M urea. The sRNAs were transferred to a positively charged membrane and UV-fixed. GFP-specific probes were generated by PCR (forward primer: 5’-AGATCCGCCACAACATCGAG, reverse primer: 5’- TTGTACAGCTCGTCCATGCC) and labelled with the DIG DNA Labelling Mix (Roche, bMannheim, Germany). After hybridisation, detection was done by incubating membranes with Anti-Digoxigenin-AP (Roche, Mannheim, Germany) followed by signal detection with the CDP-Star substrate (Roche, Mannheim, Germany) using an iBright FL1500 imaging system (Thermo Fisher Scientific, Waltham, MA, USA).

For the Western blot analysis, an aliquot of the IP eluate was separated on a 15% polyacrylamide gel by electrophoresis, transferred to a nylon membrane and incubated with a rabbit anti-HA antibody (1:1000 dilution), followed by a goat anti-rabbit secondary antibody conjugated to horseradish peroxidase (1:10000 dilution). For signal detection, the membrane was incubated with the Clarity™ Western ECL Substrate (Bio-Rad, Hercules, CA, USA), and signal was detected using the iBright FL1500 imaging system.

### sRNA Sequencing and Bioinformatic Analysis

Library preparation and sequencing were conducted at the Ramaciotti Centre for Genomics (UNSW Sydney, Australia) and the Central Analytical Research Facility (CARF) at QUT (Brisbane, Australia). Libraries were prepared using the QIASeq miRNA library kit (Qiagen, Hilden, Germany), with total RNA samples undergoing rRNA depletion using the QIAseq FastSelect kit (Qiagen, Hilden, Germany). Sequencing was done on a NovaSeq X Plus using a 10B flow cell (LEL13934) or a NextSeq 2000 using a P1 100 flow cell (Illumina, San Diego, USA). Preliminary quality checks, read mapping and coverage statistic calculation were performed as per Gauthier et al. (2026). Plots were then derived using in-house Python and R scripts.

## Acknowledgements

This work was supported by funding from the Hort Innovation project BY22011 (CON-002530). We acknowledge Queensland University of Technology (QUT) for providing subsidised access to specialised research infrastructure, including high-performance computing (HPC), and support from QUT’s Digital Research Infrastructure team.

## Notes

### Competing Interest Statement

The authors have declared no competing interest.

