## Supplementary Figures for "VIPER-seq: Protein-mediated small RNA enrichment enhances plant virus detection at low viral abundance"

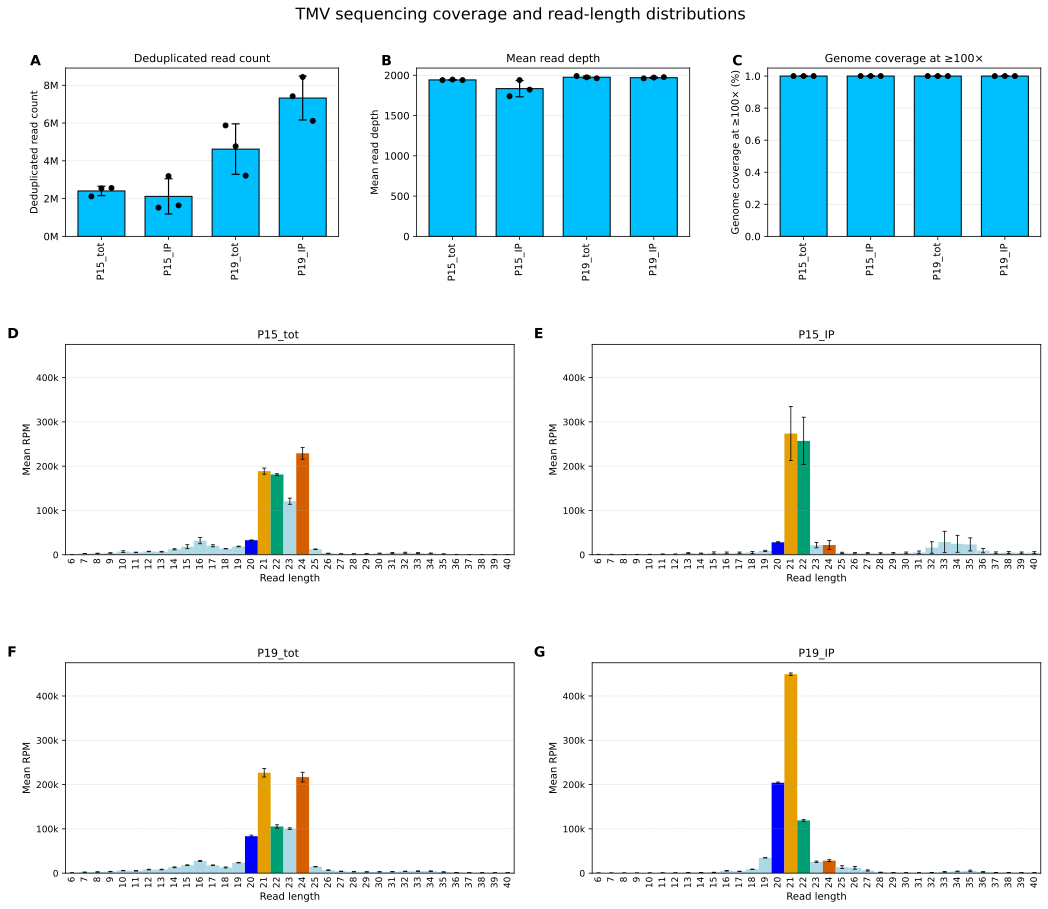

**Figure S1. TMV sequencing coverage and read-length distributions.** (A-C) Sequencing reads mapped to the TMV genome are presented as deduplicated read counts, mean read depth and genome coverage. (D-G) The size distribution of reads originating from different treatments is presented. Read lengths expected for sRNAs are highlighted with size-specific colours.

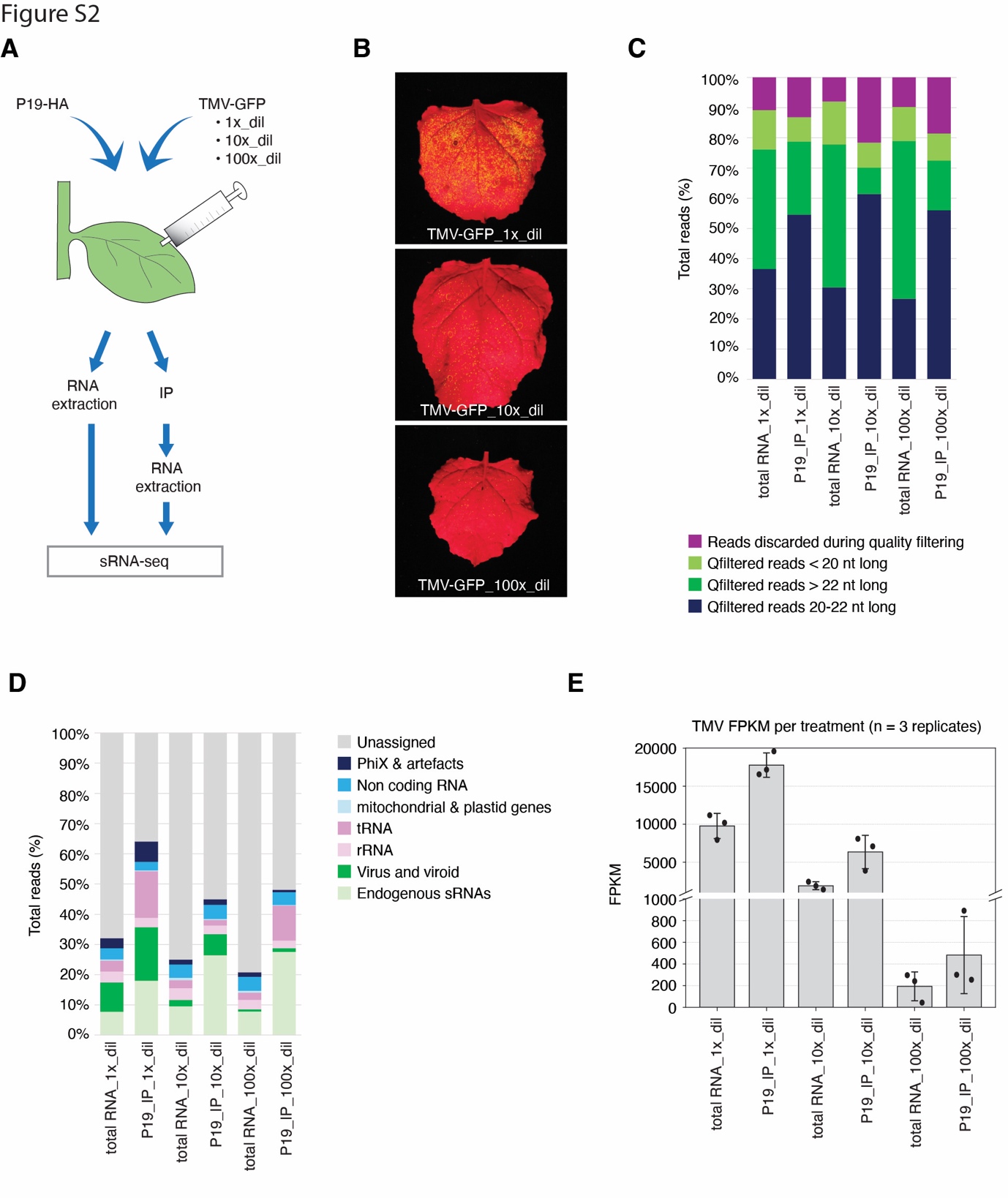

**Figure S2. Evaluation of VIPER-seq for the detection of low-titre viruses.** (A) Experimental workflow. (B) *N. benthamiana* leaves three days after infiltration with different dilutions of *A. tumefaciens* carrying the TMV-GFP infectious clone. Percentage of sequence reads by size (C) and genomic origin (D). (E) Reads mapped to the TMV genome.

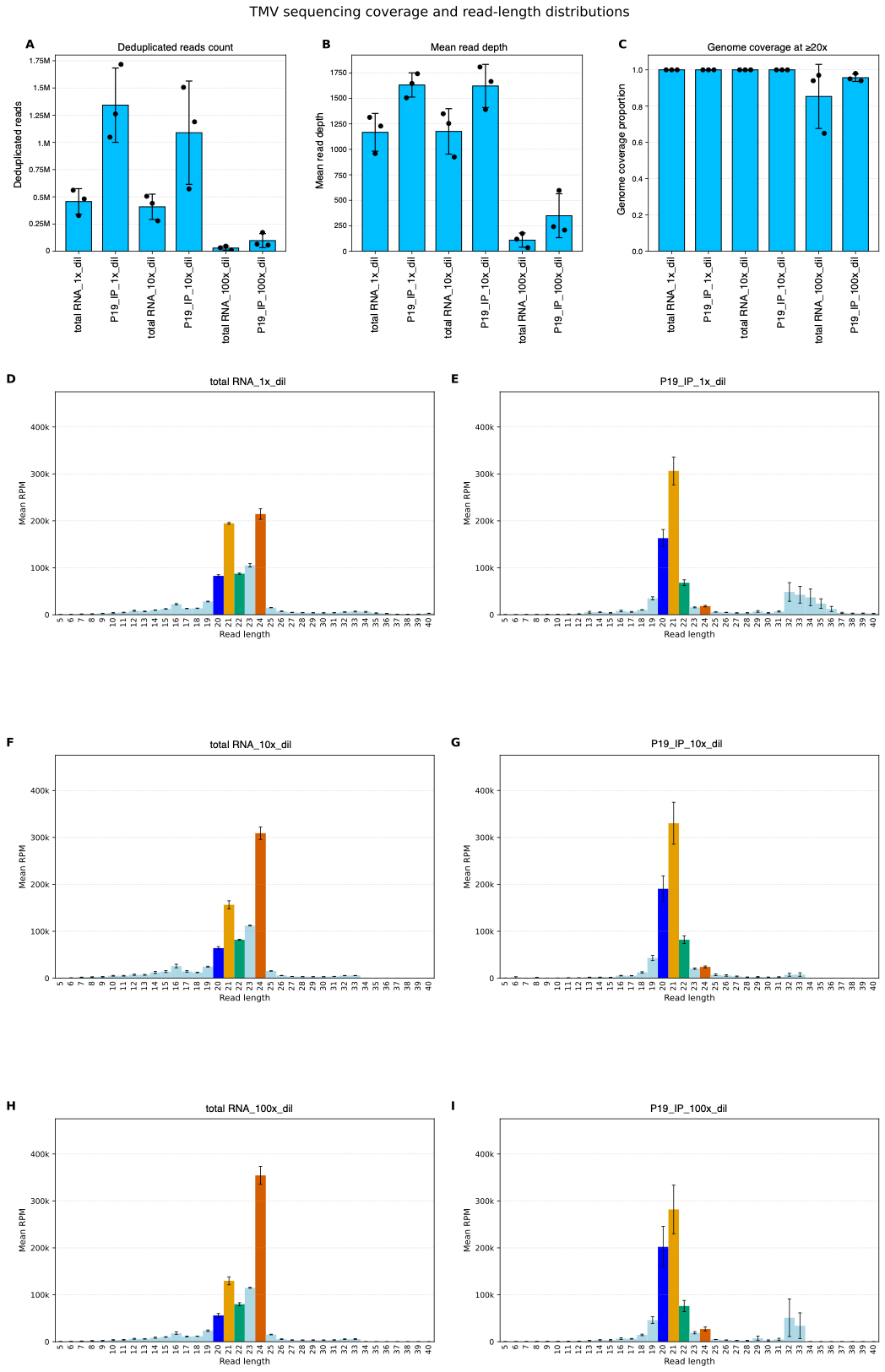

**Figure S3.** **Performance of VIPER-seq under low viral load conditions.** TMV-associated sequencing reads were assessed based on deduplicated counts (A), average coverage depth (B), and the proportion of the viral genome covered by mapped reads (C). (D-G) Read-length distributions from each treatment are shown, with colours used to distinguish size classes expected for sRNA molecules.

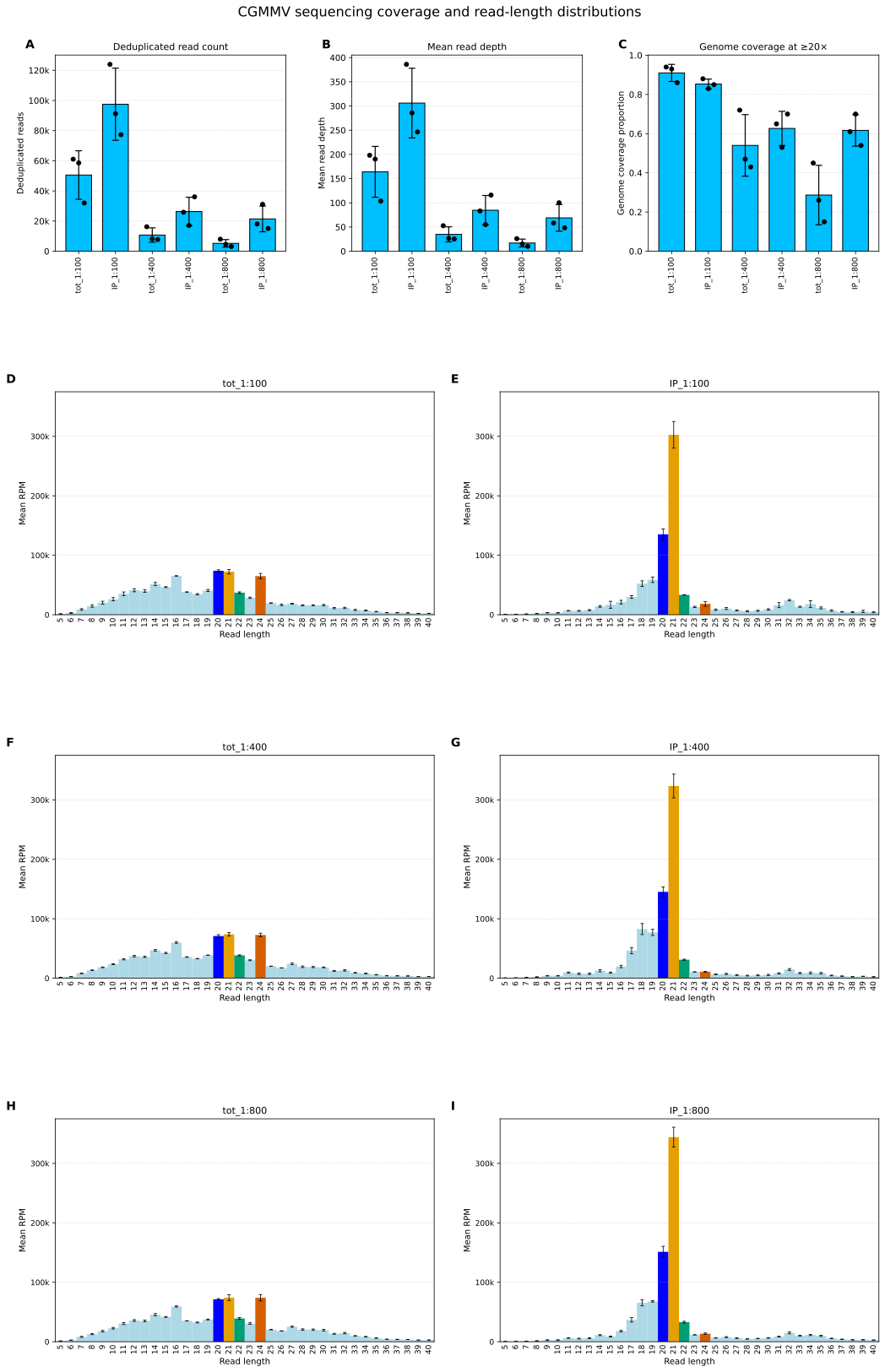

**Figure S4. Detection of low-abundance viral infection in cucumber seeds using VIPER-seq.** (A-C) Sequencing reads aligned to the CGMMV genome are presented as deduplicated read counts, average read depth, and genome coverage. (D-G) Read-length distributions for each treatment are shown. Size ranges corresponding to sRNAs are highlighted using length-specific colour coding.
